# Phylogenetics and genomic variation of two genetically distinct *Hepatocystis* clades isolated from shotgun sequencing of wild primate hosts

**DOI:** 10.1101/2024.06.21.600103

**Authors:** Paige E. Haffener, Helena D. Hopson, Ellen M. Leffler

## Abstract

*Hepatocystis* are apicomplexan parasites nested within the *Plasmodium* genus that infect primates and other vertebrates, yet few isolates have been genetically characterized. Using taxonomic classification and mapping characteristics, we searched for *Hepatocystis* infections within publicly available, blood-derived low coverage whole genome sequence (lcWGS) data from 326 wild non-human primates (NHPs) in 17 genera. We identified 30 *Hepatocystis* infections in *Chlorocebus* and *Papio* samples collected from locations in west, east, and south Africa. *Hepatocystis cytb* sequences from *Papio* hosts phylogenetically clustered with previously reported isolates from multiple NHP taxa whereas sequences from *Chlorocebus* hosts form a separate cluster, suggesting they represent a new host-specific clade of *Hepatocystis.* Additionally, there was no geographic clustering of *Hepatocystis* isolates suggesting both clades of *Hepatocystis* could be found in NHPs throughout sub-Saharan Africa. Across the genome, windows of high SNP density revealed candidate hypervariable loci including *Hepatocystis*-specific gene families possibly involved in immune evasion and genes that may be involved in adaptation to their insect vector and hepatocyte invasion. Overall, this work demonstrates how lcWGS data from wild NHPs can be leveraged to study the evolution of apicomplexan parasites and potentially test for association between host genetic variation and parasite infection.

**Author Summary:** Non-human primates are hosts to many species of *Plasmodium*, the parasites that cause malaria, and a closely related group of parasites called *Hepatocystis*. However, due to restrictions and challenges of sampling from wild populations, we lack a complete understanding of the breadth of diversity and distribution of these parasites. Here, we provide a framework for testing already-sampled populations for parasite infections using whole genome sequences derived from whole blood samples from the host. Following taxonomic classification of these sequences using a database of reference genomes, we mapped reads to candidate parasite genomes and used an unsupervised clustering algorithm including coverage metrics to further validate infection inferences. Through this approach, we identified 30 *Hepatocystis* infections from two genetically distinct clades of *Hepatocystis* in African non-human primates and described genes that may be under immune selection in each. Most importantly, the framework here can be applied to additional sequencing datasets from non-human primates and other vertebrate hosts as well as datasets from invertebrate vectors. Therefore, this approach could greatly improve our understanding of where these parasites are found, their host-specificity, and their evolutionary history. This framework may also be adapted to study evolution in other host-pathogen groups.

## Introduction

Despite the phylogenetic placement of the apicomplexan *Hepatocystis* parasites within the *Plasmodium* genus, they maintain a different genus identifier since they lack key characteristics of *Plasmodium*. Most notably, *Hepatocystis* lacks asexual replication in red blood cells, the cause of malarial disease symptoms, and are transmitted by midges of the genus *Culicoides* instead of *Anopheles* mosquitoes(1–3). In spite of these major differences, *Hepatocystis* are a sister clade to the rodent malaria parasites(1, 3, 4), which serve as model systems for human malaria(5), and maintain a similar life cycle including infection of hepatocytes followed by erythrocytes and transmission through a vector blood meal(6, 7). Also, like *Plasmodium*, *Hepatocystis* parasites infect a wide range of vertebrate hosts including non-human primates (NHPs), although notably are not thought to infect humans(4, 7). Surveys of *Hepatocystis* species diversity have been conducted in bats(8–16) and NHPs, although more limited in the latter in part because of the difficulty of obtaining blood samples from wild individuals.

In NHPs, six species of *Hepatocystis* have been described via microscopy, all in old world monkeys (OWMs)(4). In South Asia, two *Hepatocystis* species have been reported in the genus *Macaca*: *H. semnopitheci* in long-tailed and pig-tailed macaques in southern Thailand and *H. taiwanensis* in Formosan rock-macaques in Taiwan(7, 17). Genetic surveys of *cytochrome b* (*cytb*) indicate that *Hepatocystis* is prevalent in Thai macaques (44-55% infected) but the sequence data has generally not been linked to a morphologically described species(18). The remaining four morphologically described species of *Hepatocystis* – *H. kochi*, *H. bouillezi*, *H. simiae*, and *H. cercopitheci* – have been found in multiple genera of African OWMs (*Cercopithecus*, *Cercocebus*, *Chlorocebus*, *Colobus*, and *Papio*)(7, 19–22). Infection prevalence, determined by either microscopy or mtDNA sequencing, ranges from 0% to over 60% in populations of OWMs from Cameroon, Uganda, Tanzania, Kenya, and Ethiopia(20–25). *Hepatocystis* mtDNA has also been reported in both fecal and blood samples from chimpanzees (*Pan troglodytes schweinfurthii*, *Pan troglodytes ellioti*, and *Pan troglodytes troglodytes*) sampled in Uganda, Cameroon, the Democratic Republic of Congo, and Tanzania, suggesting *Hepatocystis* also infects some great apes(25). Despite infection of multiple NHP taxa in Africa, a phylogenetic tree of *Hepatocystis cytb* sequences isolated from different species of African OWMs and chimpanzees exhibits no obvious geographic clustering or host-specificity, suggesting a single generalist *Hepatocystis* species may infect diverse African NHPs(24–26). Limited sampling, both across primates and the genome hinder a deeper understanding of co-evolution between *Hepatocystis* and their NHP hosts.

Although phylogenetic analysis of *cytb* is informative, genomic sequences are necessary to robustly infer evolutionary relationships and to gain insight into patterns of genetic variation and genome evolution. The first and only *Hepatocystis* genome sequence was published in 2020 based on sequences from an infected red colobus monkey (*Piliocolobus tephrosceles*)(3). The genome assembly revealed that several loci known to be involved in *Plasmodium* mosquito stages had a high non-synonymous substitution rate relative to *Plasmodium* and that several genes important in liver stages had increased copy numbers. These hint at possible adaptations *Hepatocystis* may have evolved for utilizing *Culicoides spp*. as a vector and primarily infecting hepatocytes, respectively(3). Additionally, genes involved in erythrocytic schizogony were either found to be present with fewer copies than in *Plasmodium*, such as pir genes, or were entirely absent, including those encoding reticulocyte binding proteins(3). While these results are groundbreaking in furthering our understanding of *Hepatocystis* evolution, they remain the only genomic sequence data for *Hepatocystis* and reflect a single species.

In this work, we aim to expand on the existing phylogenetic and evolutionary analyses of *Hepatocystis* parasites by extracting *Hepatocystis* sequences from publicly available shotgun sequences of wild NHPs. We identify infections in *Chlorocebus* monkeys and *Papio cynocephalus* from Ethiopia, Kenya, and South Africa, consistent with where infections have been reported in previous studies, as well as in Zambia and The Gambia, building on our knowledge of the range of *Hepatocystis spp*. Most notably, we identify sequences from a previously undescribed, genetically distinct species of *Hepatocystis* so far only found to infect *Chlorocebus* whereas sequences from *Papio* infections clustered with the published *Hepatocystis* reference and *cytb* sequences derived from other NHP taxa. Using the sequences obtained from both *Chlorocebus*– and *Papio-*infecting *Hepatocystis*, we explore patterns of genomic variation and identify genes potentially undergoing diversifying selection in *Hepatocystis*.

## Results

### Curation of a sequence dataset from wild, non-human primates

To survey *Hepatocystis* and *Plasmodium* infections across NHP species and geographic locations, we searched the NCBI SRA database, a database of publicly available sequence reads, for shotgun sequence data from wild NHPs derived from blood samples. We identified 326 such samples from 17 different primate species originating from 18 countries across Africa, Asia, and the Caribbean (Fig 1, S1 Table). In total, 83% of samples were from African OWMs, 3% from Asian OWMs, and 14% from great apes. Most samples (76%) were collected in Africa. Of the remaining samples, 12% were collected in the Caribbean Islands and the other 12% were collected in Asia.

**Fig 1.**
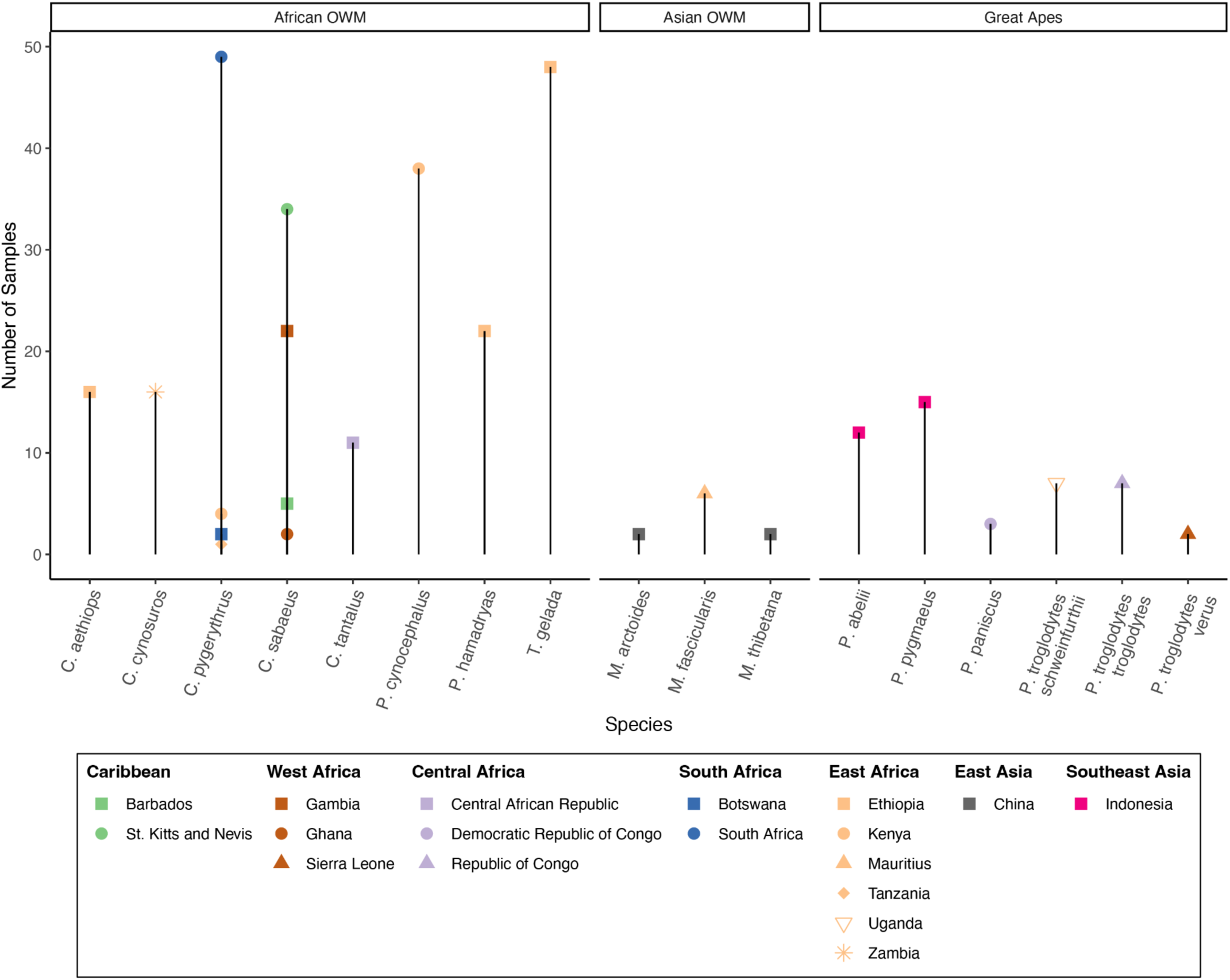
Number of samples per species in the curated dataset of blood-derived whole genome sequence data from wild non-human primates. Species are grouped by genus and taxonomic group (OWM: Old World Monkey). Each colored point represents a collection of samples (BioProject) in the curated dataset. Points are colored by geographic region and shaped by country of sampling within each region.

### *Hepatocystis* infections in African Old World Monkeys

We searched for *Hepatocystis*-derived sequences in the read data for all 326 NHP samples using Kraken2, a taxonomic classification tool that employs a kmer-based approach to classify sequence reads by comparing to a database of numerous reference genomes(27, 28). We identified 42 samples with an elevated proportion of reads classified as *Hepatocystis* from two genera: *Papio* and *Chlorocebus* (Fig 2A). For further evaluation, reads from all *Papio* and *Chlorocebus* samples were mapped to a joint primate and *Hepatocystis* reference genome (10.5281/zenodo.12209844) and we assessed the distribution of coverage across nuclear and mitochondrial (mtDNA) genomes of the parasite. K-means clustering using the ratio of mtDNA to nuclear coverage, a measure of coverage uniformity (interquartile range), and the percent of reads classified revealed two out of three clusters that corresponded to elevated values across the three predictors. We consider the 30 NHP samples in these two clusters as infected (Fig 2B, S1 Fig, and S2 Table). The infected samples include 20 individuals from four different species in the genus *Chlorocebus* and 10 *Papio cynocephalus* samples, all found in Africa (Fig 2C). The average coverage was 0.26X and 0.15X for nuclear genomes and 35X and 18X for mtDNA genomes across infected *Papio* and *Chlorocebus* samples, respectively (S2 Fig). We note that additional cases may be true infections, particularly those with elevated values for only one or two of the predictors, which tend to be samples with overall lower coverage (S2 Fig). Although we applied the same pipeline to look for *Plasmodium*, we found no clear infections supported by multiple predictors.

**Fig 2.**
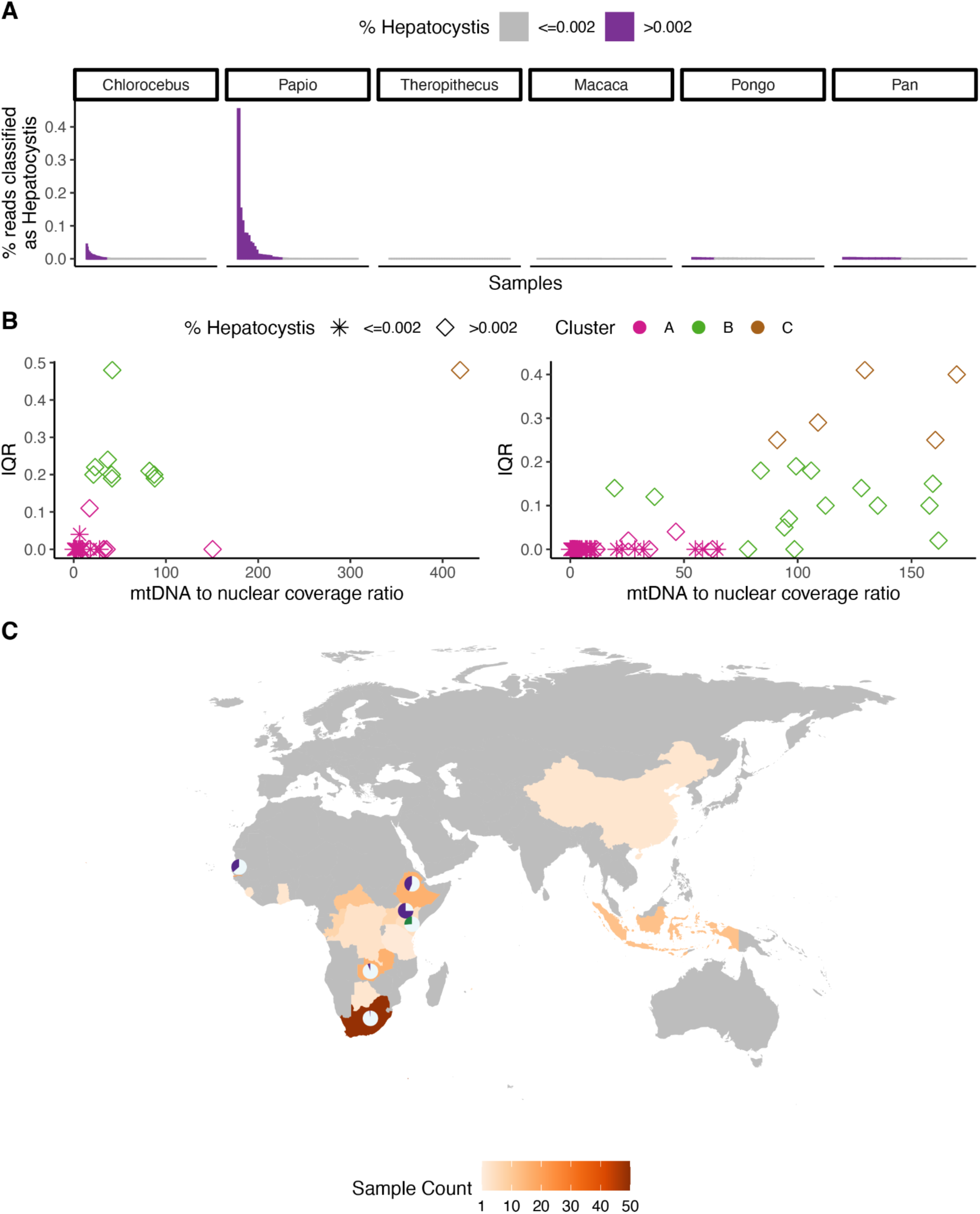
Summary of infection inference. A) Percent of reads classified as *Hepatocystis* by Kraken2 for all 326 samples. Samples with >0.002% of reads classified as *Hepatocystis* (more than 6SD above the mean for uninfected human samples) are colored in purple. Samples are grouped by genus. B) Relationship between predictor variables used in K-means clustering of samples to infer infection status of *Papio* samples (left) and *Chlorocebus* samples (right) using % reads classified as *Hepatocystis*, ratio of mtDNA to nuclear coverage and interquartile range of coverage (IQR) as predictors. Data points are shaped by the percent of reads classified by Kraken2 and colored by the cluster from the K-means algorithm using k = 3. In both pHep and cHep, clusters B and C were inferred as infected. C) Map showing sampling locations for all 326 samples. Countries with pie charts are locations where some samples were inferred as infected. In the pie charts, gray is the proportion uninfected and purple (*Chlorocebus*) and green (*Papio*) are the proportion infected. There were no samples from the countries colored gray.

### Host-species specificity in African OWM-infecting *Hepatocystis*

Next, we assessed the level of host specificity across infected samples in this dataset by assembling a *Hepatocystis cytb* sequence from each of the 30 infected samples and constructed a phylogeny. The resulting phylogeny divides the *Papio*-infecting (pHep) and *Chlorocebus*-infecting (cHep) samples into two clades with strong bootstrap support, indicating host specificity between these two primate groups at a genus level (Fig 3A). Within the cHep clade, there are three clusters: one containing six sequences derived from *C. aethiops* samples, another with five sequences from *C. sabaeus* samples, and another with sequences from one *C. pygerythrus* and one *C. cynosuros* sample. Branch lengths were short (< 0.006) but suggest potential substructure at the species level. In the pHep clade, the *cytb* sequences from *Papio* samples clusters with the sequence found in the *Hepatocystis* reference genome, derived from a *Piliocolobus* monkey. The branch length of 0.032 between the pHep and cHep clades suggests they are likely to represent two *Hepatocystis* species(26).

**Fig 3.**
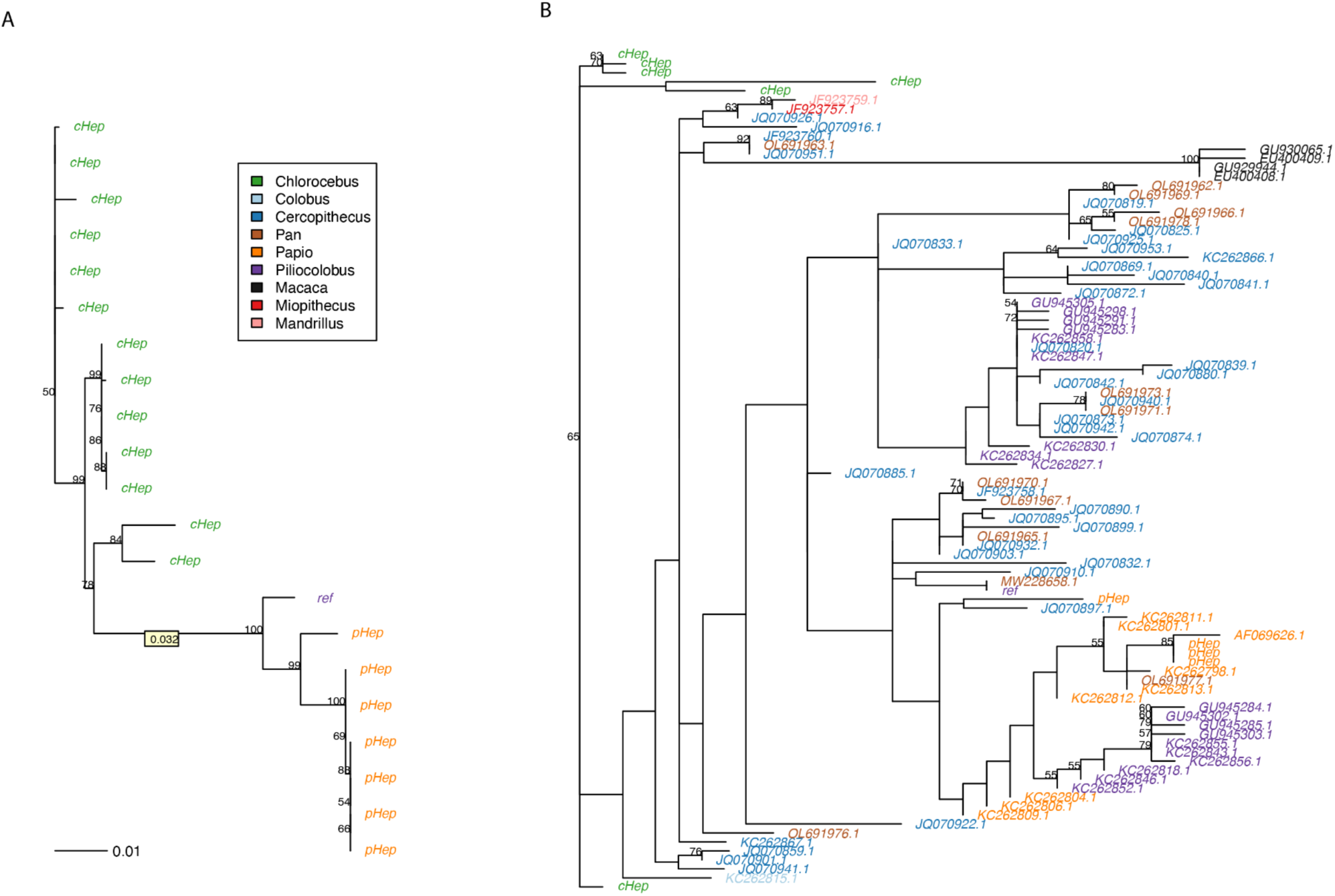
Phylogenetic trees inferred with PhyML of the unique *cytochrome b (cytb)* sequences. The trees are constructed from the dataset presented here together with A) the *Hepatocystis* reference sequence (alignment length=1.1kb) and B) additional unique publicly available *Hepatocystis* sequences from various NHPs identified via PCR (689bp). Tip labels are colored by the host genus as given in the legend. Bootstrap values >50% are shown. The branch length between the *Chlorocebus*-infecting (cHep) and *Papio*-infecting (pHep) isolates is highlighted in a yellow box in panel A.

To put this into a broader evolutionary context, we combined our assembled *cytb* sequences and the reference sequence with 86 publicly available, unique *Hepatocystis cytb* sequences that have been identified in a range of NHPs(17, 18, 23–25). The resulting phylogeny further supports a distinct, host-specific cHep clade (Fig 3B). In contrast, pHep sequences cluster with multiple *Piliocolobus*-derived sequences and with little evidence for host-specificity across this larger phylogeny, suggesting this species of *Hepatocystis* may be a generalist across African NHP hosts. Additionally, pHep sequences appear more closely related to the clade of Asian NHP *Hepatocystis* sequences than they are to the cHep clade although we note that the placement of Asian NHP infecting sequences within this phylogeny has low bootstrap support.

### Genomic variation in *Hepatocystis*

Although the average coverage across the nuclear genome was low (mean 0.26X in pHep and 0.15X in cHep), we were able to identify genomic variation by applying a probabilistic approach accounting for genotype uncertainty. Throughout the genome we identified 20,955 SNPs in pHep and 41,099 SNPs in cHep (10.5281/zenodo.12209844). This allowed us to look at SNP density across the genome to identify hypervariable loci that may be involved in immune evasion in *Hepatocystis* (Fig 4). We first tested our ability to recover hypervariable regions from low coverage data by applying the same pipeline to calculate SNP density in a *P. falciparum* dataset down sampled to 30X (378,737 SNPs), 1X (55,759 SNPs), 0.5X (26,281 SNPs) and 0.1X (1,117 SNPs) coverages. We used precision recall curves to determine a threshold for considering windows of high SNP density (S3 Fig). Using 1kb bins in the 30X data with a SNP density greater than 2 standard deviations from the mean as the truth set, both 1X and 0.5X datasets had high precision (100% for the top 10 bins and >96% for the top 50 bins) and overlapped genes known to be hypervariable in *P. falciparum* (S3 Table). The 0.1X dataset did not perform as well, but the number of SNPs in this dataset for *P. falciparum* was extremely low, likely due to the much lower genetic diversity (∼10x lower) than most other *Plasmodium*(29). The number of SNPs in our dataset (41,099 for cHep and 20,955 for pHep) more closely match 0.5X and 1X for P. falciparum. We therefore chose to consider the top 50 bins in our dataset as candidate hypervariable regions.

**Fig 4.**
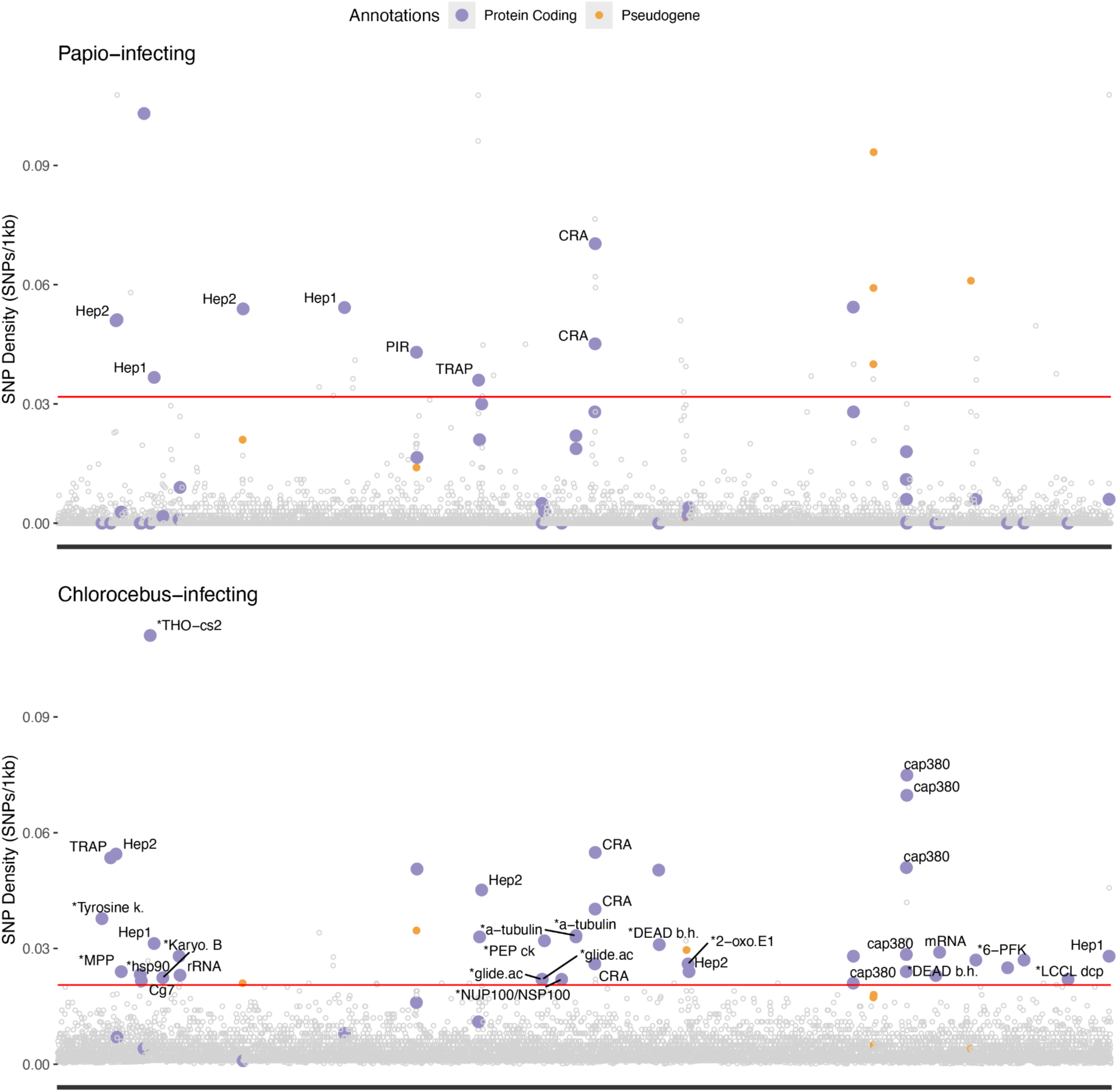
SNP density in pHEP (top) and cHEP (bottom) in 1kb bins. In both plots, the solid red line represents the threshold for the 50 windows with the highest SNP densities. The top 50 points are annotated as follows: purple points indicate bins that overlap protein coding genes and are labeled with the gene name, if one has been assigned (* indicates a shortened name, with full names given in S2 Table). Orange points overlap pseudogenes. Gray points above the line do not overlap a gene. Points in the top 50 of either pHep or cHEP are colored in both plots. Points not in the top 50 in either plot are gray.

Protein-coding genes (and not pseudogene members) from the *Hepatocystis-*specific *Hep1* and *Hep2* gene families were among the most hypervariable in both pHep and cHep (Fig 4; S4 and S5 Tables). Both gene families were first identified in the *Hepatocystis* reference assembly and are unique to *Hepatocystis* with unknown function(3). Of the 16 genes in the *Hep1* gene family, we identified three as hypervariable in either pHep or cHep and one (HEP_00519600) as hypervariable in both. Of the 10 genes in the *Hep2* gene family, two were hypervariable in either pHep or cHep and one was hypervariable in both (HEP_00480100).

Multiple 1kb windows spanning the entire circumsporozoite-related antigen gene (CRA; HEP_00212100 in *Hepatocystis*) were hypervariable in both pHep and cHep. CRA is conserved across *Plasmodium* and is involved in hepatocyte invasion(30, 31). Two genes expressed in stages within the vector, thrombospondin-related anonymous protein (TRAP) and the oocyst capsule protein (Cap380), were found to be among the most hypervariable as well. The TRAP gene in *Plasmodium* is involved in salivary gland infection within the mosquito(32). Unlike *Plasmodium*, *Hepatocystis* was found to have six copies of TRAP in the reference genome(3). We identified one TRAP gene as hypervariable in pHep (HEP_00163800) and another in cHep (HEP_00475600). Cap380 is essential for survival and development of *Plasmodium* oocysts into sporozoites within the mosquito(*33*). Several bins with high SNP density spanned the Cap380 gene region in cHep, but not in pHep. Although conserved across *Plasmodium* species(31, 33, 34), Cap380, CRA, and TRAP genes, were not found to be hypervariable in our analysis of *P. falciparum*. In fact, in the 30X *P. falciparum* dataset, there were no hypervariable genes that were also hypervariable in *Hepatocystis*. Lastly, the most hypervariable bin in cHep overlapped with the THO complex subunit 2 gene (THO2, HEP_00514500; Fig 4 and S5 Table). Across eukaryotes, THO2 is part of the highly conserved TREX complex involved in mRNA export, but in *Plasmodium* some of the TREX complex subunits are absent, and THO2 lacks a conserved domain(35).

### No evidence for association between *ACKR1* genetic variation and *Hepatocystis* infection in *Chlorocebus*

In humans, a SNP in the GATA-1 transcription factor binding region upstream of the *ACKR1* gene is associated with protection against *P. vivax* (rs2814778, –67 T>C)(36, 37). Similarly, it has been suggested that a SNP in the 5’UTR of the *ACKR1* gene in *Papio cynocephalus* may reduce susceptibility to *Hepatocystis* infection (A>G 359bp upstream of the transcription start site)(38). Using the inferred infection status and host genetic variation data, we sought to test for association between *ACKR1* variation and *Hepatocystis* infection in this dataset. Since the *Papio* samples had a lower average host coverage than the *Chlorocebus* samples (1.5X vs 5X), we tested for association in *Chlorocebus* using the publicly available variant calls(39). When considering all *Chlorocebus* samples, we identified five nonsynonymous SNPs within the *ACKR1* gene, including two within extracellular domains, and eight within 1,000 bp upstream of the *ACKR1* 5’UTR that could potentially be involved in regulation of *ACKR1* and encompass the regulatory region where associations have been reported. However, when only considering SNPs segregating in countries where infections were present, none were significantly associated with *Hepatocystis* infection (S6 Table). We observed four missense and two synonymous variants as well as one intronic and one upstream variant that were present in uninfected samples and absent from infected samples, potentially consistent with a protective effect. However, larger samples will be required for sufficient power to evaluate this (S6 Table).

## Discussion

In this study, we analyzed publicly available whole genome sequence data to survey the distribution of *Hepatocystis* in 326 wild NHPs from Africa and Asia. Notably, of the 17 species of NHPs we were able to include, five do not have published surveys for *Plasmodium* or *Plasmodium-like* infections(40). These species are *Pongo abelii*, *Theropithecus gelada*, *Macaca thibetana*, *Chlorocebus cynosuros*, and *C. tantalus*. However, we note that it is possible that previous surveys were negative and the findings were not published. Alternatively, they could have been surveyed under different species names. For instance, many *Chlorocebus* species were considered as *Cercopithecus aethiops* and various subspecies until 1996(41, 42). Similarly, *P. abelii* was not classified as a species until the mid-2000s(43). Thus, if we assume they have not been previously sampled, *C. cynosuros*, could be a possible new host of *Hepatocystis* as we inferred an infection in this species. This also suggests *Hepatocystis* is circulating in the Kafue region of Zambia where this individual was sampled from. We additionally report *Hepatocystis* infections in The Gambia for the first time, in nine of 20 *C. sabaeus* samples, highlighting the utility of surveying parasites in wild NHP whole genome sequence data as it becomes available to continue improving our understanding of the distribution of *Hepatocystis* or other blood-borne pathogens.

We also identify infections in species that are known hosts of *Hepatocystis*: *Papio cynocephalus*, *Chlorocebus aethiops, and Chlorocebus pygerythrus*(7, 20–22, 44). In this dataset, these samples were collected from locations in Ethiopia, Kenya, and South Africa. Ethiopia has the highest geographic representation, comprising 25% of all samples in the dataset including *Papio hamadryas* (n = 20), *C. aethiops* (n = 16), and *Theropithecus gelada* (n = 48)(39, 45). Of the Ethiopian samples, *C. aethiops* was the only species inferred to be infected with *Hepatocystis* in our analysis, consistent with previous surveys of the *Chlorocebus* genus and *P. hamadryas* in Ethiopia(20, 22). The higher altitude environment at which *T. gelada* lives may drive the absence of *Hepatocystis*(45), but other environmental factors, species-barriers, or vector preferences could also be responsible especially given the absence of *Hepatocystis* in *P. hamadryas* as well. Additional sampling would be needed to confirm absence of *Hepatocystis* in *Theropithecus* and *Papio* species in Ethiopia.

In Kenya, both *P. cynocephalus* and *C. pygerythrus* carried *Hepatocystis* infections. Although these two species are found in close geographic proximity, our phylogenetic analysis indicates genetically distinct parasites in each genus, which we refer to as pHep and cHep since we do not have morphological data for species assignment. Notably, in South Africa, Ethiopia, Kenya, Zambia, and The Gambia, four different species of *Chlorocebus* were infected with genetically similar parasites, suggesting they represent a *Chlorocebus*-specific clade of *Hepatocystis* (cHep). In contrast, the *Hepatocystis isolates* found in *P. cynocephalus* (pHep) in this study phylogenetically clustered with previously reported *Hepatocystis* isolates from *Mandrillus*, *Miopithecus*, *Cercopithecus*, *Piliocolobus*, *Colobus*, and *Pan*, therefore spanning multiple OWM and great ape species(23–25). Neither cHep nor pHep exhibit geographic specificity, as the previously reported isolates of *Hepatocystis* that clustered with pHep were collected from NHPs in Cameroon and Uganda.

Despite these differences in host-specificity and geographic distribution, in both cHep and pHep we find evidence for diversifying selection on genes in two *Hepatocystis-*unique gene families, Hep1 and Hep2. Although their function remains unknown, the expansion of both families and high SNP density suggest they may be involved in immune evasion. Several Hep1 and Hep2 genes are highly expressed in blood stages, although unlike *Plasmodium*, *Hepatocystis* does not undergo asexual replication in the blood stage(3). Despite lacking this stage and the associated pathogenic outcomes, *Hepatocystis* may utilize the Hep1 and Hep2 gene families in a similar strategy for immune evasion as other hypervariable gene families across *Plasmodium* (e.g., var, SICAvar, pir). The TRAP gene family has also expanded uniquely in *Hepatocystis*, with one of the six genes in the family appearing as hypervariable in pHep and another in cHep. In *Plasmodium*, TRAP plays a role in infection of salivary glands in the mosquito and hepatocytes, suggesting potentially increased conflict between *Hepatocystis* and either the liver or vector stage. Sporozoites are larger in *Hepatocystis* than *Plasmodium* and have a longer prepatent period in the liver where it is thought to also be able to sustain chronic infection(*2*). CRA, a single copy gene with a role in hepatocyte invasion, also has high SNP density in both pHep and vHep. Finally, the gene cap380, which is involved in oocyst development into sporozoites within the insect vector, shows the highest SNP density in the genome in cHep but is not hypervariable in pHep, possibly resulting from interaction with a different *Culicoides* vector species. However, we note that differences between high SNP density windows in cHep compared to vHep could also be due to limited sensitivity in low coverage data.

Paired with host variation data, infection inference allows for the potential discovery of SNPs associated with *Hepatocystis* infection. Although the current sample size and host genomic coverage limit this application, we demonstrate this potential by testing for association between *Hepatocystis* infection and variation in the *ACKR1* gene, which has previously been associated with *Hepatocystis* infection in baboons(38). We identified eight alleles present only in uninfected *Chlorocebus* samples, but none were significantly associated with infection status given the small sample size. Additionally, two of these are missense variants in extracellular regions, although not within the region where *P. vivax* is thought to bind to the receptor in humans(46). Future studies with larger samples and in baboons could shed light on whether there is consistent evidence for association between *Hepatocystis* and *ACKR1* variation in NHPs, paralleling the association between *ACKR1* and *P. vivax* in humans. However, it remains unknown whether *Hepatocystis* uses ACKR1 as a receptor or whether *Hepatocystis* has been a strong selective pressure given the milder disease manifestation.

In this analysis, we did not find any confidently inferred *Plasmodium* infections upon combining Kraken2 read classification and coverage across the nuclear and mitochondrial genomes. This is likely a sampling bias rather than any inherent difference in detectability using our pipeline, as African great apes, hosts of the Laverania clade of *Plasmodium*, and macaques, hosts of at least four species of *Plasmodium*, were underrepresented in our study and the remainder of African NHPs included are thought not to carry natural infections of *Plasmodium*. Increased sampling from wild NHP populations would therefore be an opportunity to identify *Plasmodium* infections as well. We were able to infer *Hepatocystis* infections in samples with as low as 0.4X host genome coverage, suggesting this approach is applicable even to low-coverage NHP sequencing projects as they become available. Nonetheless, lower coverage of both host and parasite genomes limits the detection of more complex variant types and the application of tests for selection. We also attempted to run a similar pipeline on samples from fecal and non-blood tissue samples, which would broaden the opportunities for parasite identification to non-invasively collected samples. However, we found much noisier read classification in these sample types and additional methods development may be required for this application.

Overall, we demonstrate that shotgun sequences derived from whole blood can be used to identify apicomplexan parasites and their distribution across NHP populations. We describe a putatively novel *Hepatocystis* species, cHep, emphasizing that there is still much to uncover about the diversity of *Plasmodium* and *Plasmodium*-like parasites in NHPs. Since many limitations exist regarding sampling from wild NHP populations, implementation of this approach will improve our ability to study parasite evolution and host-parasite relationships as sequencing data from more wild populations becomes available(47, 48). Notably, this approach can be adapted to study infections and associations in other vertebrate hosts of *Plasmodium*, invertebrate vectors, or even expanded to other host-parasite systems that may be difficult to sample in the wild.

## Materials and Methods

### Data Curation

We searched the NCBI SRA database (https://www.ncbi.nlm.nih.gov/sra) to compile a dataset of publicly available, wild, non-human primate sequences from blood samples. Similar to Hernandez et al. 2020(49), we performed a search for “(primate OR primates) AND (genome or genomic) NOT (Homo sapiens)”. We also did a specific search for each recognized non-human primate species, as listed in S1 Table from Hernandez et al. 2020, e.g., “Macaca fascicularis AND whole genome”. Search results were then filtered to only include samples described as “wild” and where a geographic location was provided. Paired FASTQ files for the 326 samples meeting these criteria were downloaded using sra-toolkit(50) (S1 Table). The search covered sequences available as of August 31^st^, 2022.

### Taxonomic classification with Kraken2

We created a custom Kraken2 database using the steps listed in the online manual section 9 (https://github.com/DerrickWood/kraken2/wiki/Manual) and included the following libraries: Protozoa, Archaea, Bacteria, and Virus(27, 28). The Protozoa library contains 13 *Plasmodium* reference genomes. Because of contamination from adapter, vector, or primer sequences that can sometimes be found in sequence data and reference genomes, we also included the decontamination library, Univec(51). Three primate reference genomes were also included to assess the number of reads classified as primate origin for each sample, allowing us to later calculate the proportion of reads inferred as each parasite relative to primate, and to prevent any primate sequences from being incorrectly classified as parasite sequences. The primate genomes included were:

- Human GRCh38 (https://ftp.ncbi.nlm.nih.gov/genomes/all/GCF/000/001/405/GCF_000001405.40_ GRCh38.p14/)
- Rhesus Macaque Mmul10 (https://ftp.ncbi.nlm.nih.gov/genomes/all/GCF/003/339/765/GCF_003339765.1_ Mmul_10/)
- Gray Mouse Lemur Mmur_3.0 (https://ftp.ncbi.nlm.nih.gov/genomes/all/GCF/000/165/445/GCF_000165445.2_ Mmur_3.0/)

We additionally added the Hepatocystis reference genome to the Kraken2 database (https://ftp.ncbi.nlm.nih.gov/genomes/all/GCA/902/459/845/GCA_902459845.2_HEP1/).

We then ran Kraken2 on all sequence runs from the 326 samples identified in the data search as well as 119 paired FASTQ files from the French HGDP population (corresponding to 10 samples in total)(52). The French HGDP population served as a negative control since this population is not expected to be infected by *Plasmodium* or *Hepatocystis* parasites. For samples with multiple runs, each set of paired FASTQ files were input separately to Kraken2 and then combined to the sample level for downstream analysis. The ‘--report’ flag was included to generate the standard Kraken2 output as well as a summarized classification file for downstream analysis.

### Inference of individual infection status

Three criteria were used for inferring infection status: the proportion of reads classified as parasite, the ratio of mtDNA to nuclear coverage (expected to be >1 in true infections), and the interquartile range of coverage (a measure of coverage uniformity).

For each sample, we calculated the proportion of reads classified from each *Plasmodium* and *Hepatocystis* species individually, by dividing the number of reads classified as the specific parasite species by the sum of reads classified as any primate taxon and the number of reads classified as that parasite species, in other words:

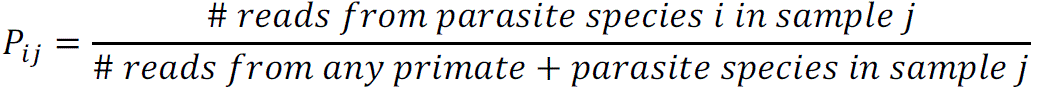

As a guide, we considered samples where the proportion of reads classified as a parasite species was more than six standard deviations greater than observed in the HGDP French population as potential infections. To further confirm positive infections in these samples, we then aligned the sequence runs to a joint primate and parasite reference genome for the specific primate and parasite pair using the closest available reference genome(53). Coverage of the parasite genome was calculated in 1kb bins using mosdepth(54) in order to determine mean mitochondrial and nuclear coverage as well as the interquartile range of coverage. Using the ratio of mtDNA to nuclear DNA, the interquartile range of coverage across the nuclear genome, and the percent classified by Kraken2, we used k-means clustering in R(55) (k = 3) to identify samples with high values in all three statistics consistent with a true infection.

### Phylogenetic analysis of *cytochrome b*

To assemble the *cytochrome b* (*cytb*) sequence from each infected sample, we extracted read pairs that mapped to a 1.1kb region of the *Hepatocystis cytb* gene (HEP_MIT003, LR699572.1:5426-6550). FASTQ files containing the extracted reads were input for assembly with SPAdes(56) with a requirement of at least two reads covering each position (coverage >= 2X), We used the reference sequence as input for the ‘--trusted-contigs’ flag to improve assembly of the gene region.

We first compared cytb sequences from the 42 infected samples. After removing duplicate sequences, 20 unique sequences were aligned using MUSCLE(57) to obtain a multiple sequence alignment in PHYLIP format. The PHYLIP file was used as input for PhyML to infer a phylogenetic tree with 100 bootstrap replicates(58). To compare the sequences with previously published phylogenies, we downloaded all available Hepatocystis cytb sequences from three published studies(17, 18, 23–25) and one unpublished study (NCBI Popset 293411005) from the NCBI nucleotide database. This dataset was reduced by removing duplicates and then only including sequences infecting greater spot-nosed monkeys that represented unique haplotypes as described by Ayouba et al.(23). Of the two macaque-infecting datasets, we used just two macaque-infecting sequences from each(17, 18) given all the sequences exhibited very high sequence similarity. This resulted in 86 additional sequences that were combined with 10 unique sequences we assembled with SPAdes. The 96 sequences were aligned with MUSCLE and trimmed using GBLOCKS, included in SeaView(59), to address the differences in regions sequenced across studies. This resulted in a 689bp sequence alignment with 96 unique sequences. As before, we inferred a phylogenetic tree with 100 bootstrap replicates in PhyML.

### Analysis of genome-wide genetic variation in *Hepatocystis*

Given that per sample coverage of the *Hepatocystis* nuclear genome was low (average coverage 0.32X), we used ANGSD(60) to calculate summaries of genetic variation in a probabilistic framework using genotype likelihoods. We called SNPs using the SAMtools model (‘-GL1’) for estimating genotype likelihoods and allele frequencies for the major and minor alleles (‘-doMajorMinor 1’, ‘-doMAF 2’), and only kept SNPs with a significant P-value (p < 1e-06). The number of SNPs per 1kb window was calculated for biallelic variants. To assess how well we could capture genomic variation with the low coverage data, we also applied the same ANGSD pipeline to 25 publicly available high coverage *P. falciparum* genomes(61) subsampled with SAMtools(62) to compare a standard high coverage of 30X with two low coverages: 1X and 0.5X. We considered the top 1100 windows (> 2SD from the mean SNP density) as true positives in the 30X data. We then plotted precision and recall using cut-offs for true positives in the low coverage datasets ranging from 10 to 100 in increments of 10 (S3 Fig).

### Association of ACKR1 variation with infection status in Chlorocebus

To transfer annotations from the orthologous regions of the human gene model, we used BLAT(63) from the UCSC genome browser with the sequence of the canonical human *ACKR1* transcript (ENST00000368122.4/NM_002036.4) as the query and the *Chlorocebus sabeus* reference genome (chlSab2). We then annotated the location of the GATA1 binding region and the human SNP within it (rs2814778) encoding the Fy^ES^ (Duffy null) allele and the SNP determining the Duffy A/B blood group polymorphism (rs12075) onto the corresponding location in the *C. sabeus* sequence.

We extracted variant calls for *Chlorocebus* samples included in our dataset from the publicly available VCF file of biallelic SNPs in a 2.6 kb region (chr20:4742279 – 4744887) containing the *ACKR1* ortholog(39). Variant annotations were added with SnpEff(64). There were 46 SNPs in this region among the 20 infected and 143 uninfected samples. The data was filtered to only include SNPs with minor allele count > 5, resulting in 24 SNPs. Samples from countries without any infections were excluded from association testing, resulting in 20 infected and 87 uninfected individuals from four countries (S6 Table). The dataset of SNPs within samples from countries with infections was further filtered to only include polymorphic sites with a minor allele count > 5, leaving 20 SNPs. Logistic regression was used to test for association between infection status and genotypes at each of the 19 SNPs under an additive model in R v4.2.3 using the lme4 (1.1-30) package. Country was included as a random effect since each country had only a single species.

## Data Availability

All non-human primate datasets used in this analysis were publicly availably NCBI BioProjects, and accession numbers can be found in S1 Table. BAM files of reads mapped to *Hepatocystis* as well as major and minor allele calls for cHEP and pHEP are available at zenodo: 10.5281/zenodo.12209844.

## Acknowledgements

We gratefully acknowledge the support and resources from the Center for High Performance Computing at the University of Utah, especially Brett A. Milash who helped to set up the initial Kraken2 database used in this analysis. We would also like to thank Nels Elde, Sarah Bush, Aaron Quinlan, and Timothy Webster from the University of Utah for their guidance and feedback on this work.

## Supporting Information

**S1 Figure. Relationship between predictor variables used in K-means clustering of samples to infer infection status for A) Papio-infecting *Hepatocystis* and B) *Chlorocebus-infecting Hepatocystis***. Data points are shaped by the percent of reads classified by Kraken2 and colored by the cluster from the K-means algorithm using k = 3.

**S2 Figure. Bar plots comparing coverage across clusters created by k-means clustering for A)pHep and B)cHep.** Bars are colored by the percent of reads classified as *Hepatocystis* from Kraken2 output and are grouped by cluster. In both pHep and cHep, clusters B and C were inferred as infected. The rows from top to bottom are: mean mitochondrial coverage, mean nuclear coverage, and percent of reads classified.

**S3 Figure. Precision recall curve for SNP densities in the *P. falciparum* dataset using hypervariable bins in the 30X data as truth sets, defined as bins with SNP density greater than 2 standard deviations (1 SD = 0.035) from the mean (mean = 0.017)**. We then used 10 thresholds for classification as hypervariable in the low coverage data. These ranged from 10 to 100 bins in increasing in increments of 10, shown as points on the plot. Points shaped as starts represent the threshold of 50 bins.

**S1 Table. Sample information for each BioProject in the curated dataset including species, sampling locations, tissue type, number of individuals, NCBI BioProject ID, and the study DOI or Grant ID**.

**S2 Table. The total number of samples and the number of *Hepatocystis*-infected samples for each species by sampling location**.

**S3 Table. Table of hypervariable gene families in *P. falciparum*.** From left to right, the columns are: gene ID, gene annotation, and the number of times each gene is determined to be hypervariable (has overlapping 1kb bin). These results are hypervariable bins in the 1X *P. falciparum* dataset that are also hypervariable in the 30X dataset.

**S4 Table. Genomic coordinates and gene content of the 50 most hypervariable 1kb bins identified in pHep**.

**S5 Table. Genomic coordinates and gene content of the 50 most hypervariable 1kb bins identified in cHep.**

**S6 Table. Logistic regression results (odds ratio, p-value and 95% confidence intervals) for the *Chlorocebus* variants segregating in samples from countries where infections were inferred**. Allele annotations, frequencies, and counts are included.

