## Supplementary figures and images for "Phylogenetics and genomic variation of two genetically distinct *Hepatocystis* clades isolated from shotgun sequencing of wild primate hosts"

### Supplemental Figures

**Supplemental Figures**

**S1 Figure**

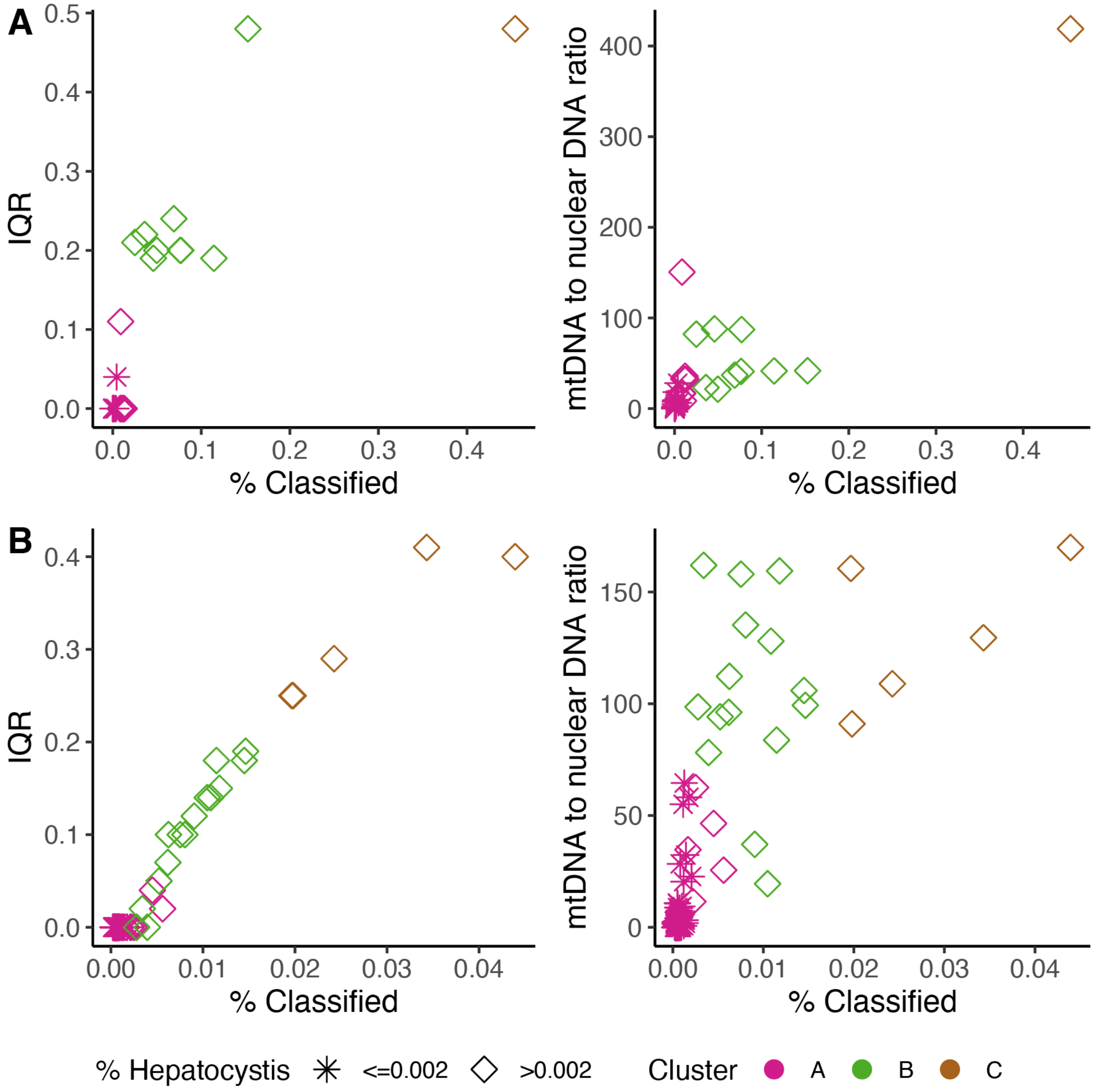

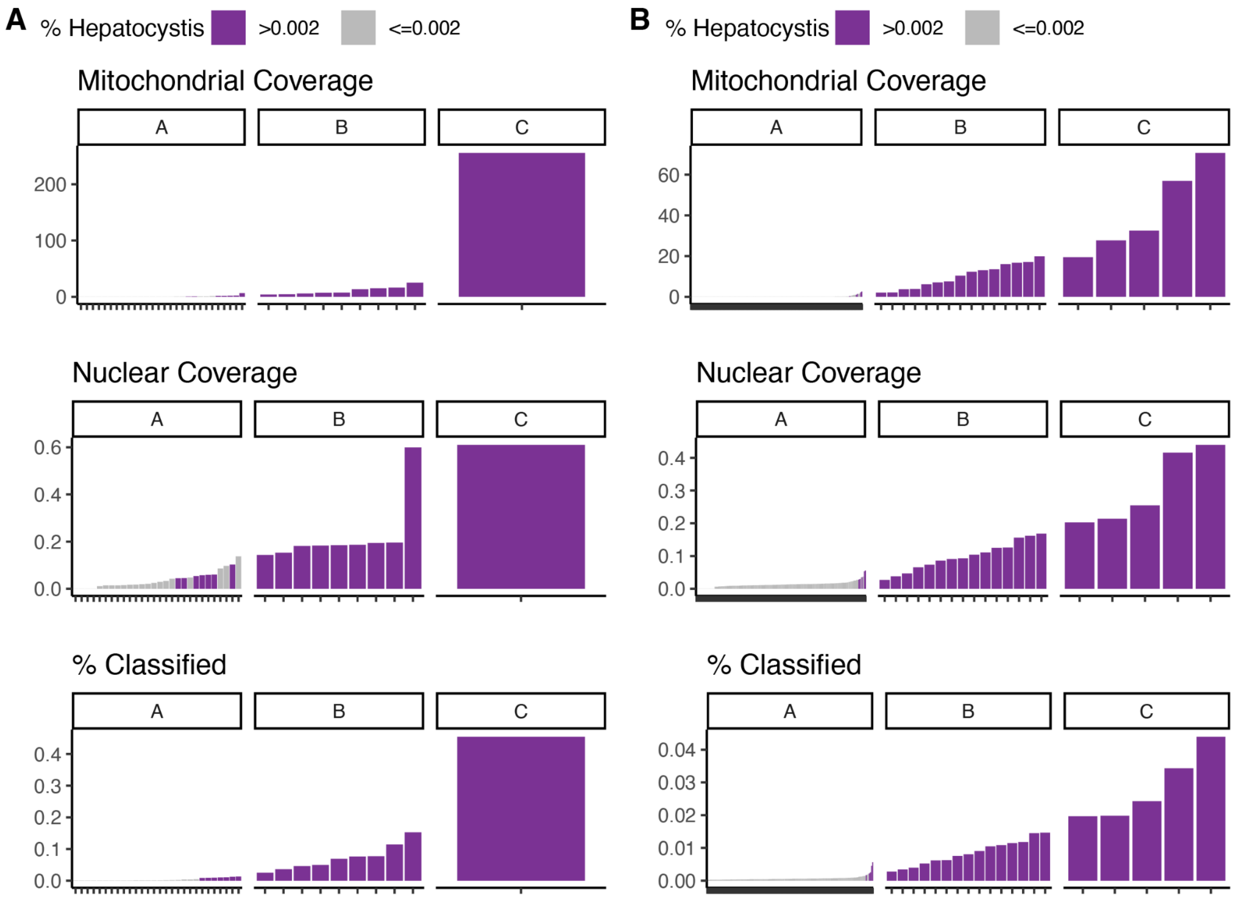

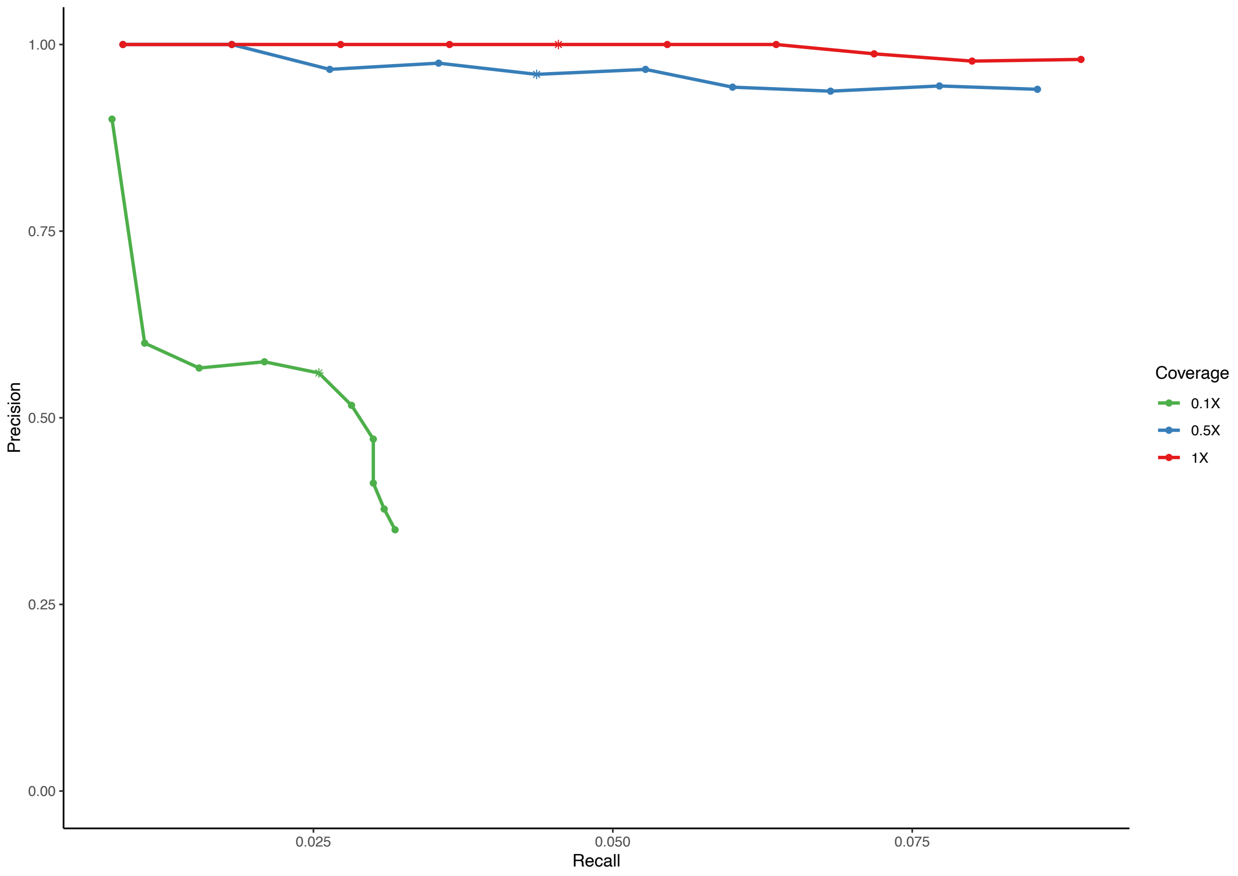
